# Using time-series remote sensing to identify and track individual bird nests at large scales

**DOI:** 10.1101/2025.02.21.639546

**Authors:** S.K. Morgan Ernest, Lindsey A. Garner, Ben G. Weinstein, Peter Frederick, Henry Senyondo, Glenda M. Yenni, Ethan P. White

## Abstract

The challenges of monitoring wildlife often limits the scales and intensity of the data that can be collected. New technologies - such as remote sensing using unoccupied aircraft systems (UAS) - can collect information more quickly, over larger areas, and more frequently than is feasible using ground-based methods. While airborne imaging is increasingly used to produce data on the location and counts of individuals, its ability to produce individual-based demographic information is less explored. Repeat airborne imagery to generate an imagery time-series provides the potential to track individuals over time to collect information beyond one-off counts, but doing so necessitates automated approaches to handle the resulting high-frequency large-spatial scale imagery. We develop an automated time-series remote sensing approach to identifying wading bird nests in the Everglades ecosystem of Florida, USA to explore the feasibility and challenges of conducting time-series based remote sensing on mobile animals at large spatial scales. We combine a computer vision model for detecting birds in weekly UAS imagery of colonies with biology-informed algorithmic rules to generate an automated approach that identifies likely nests. Comparing the performance of these automated approaches to human assessment of the same imagery shows that our primary approach identifies nests with comparable performance to human photo assessment, and that a secondary approach designed to find quick-fail nests resulted in high false positive rates. We also assessed the ability of both human photo assessment and our primary algorithm to find ground-verified nests in UAS imagery and again found comparable performance, with the exception of nests that fail quickly. Our results show that automating nest detection, a key first step towards estimating nest success, is possible in complex environments like the Everglades and we discuss a number of challenges and possible uses for these types of approaches.

## Introduction

Ecology has waited expectantly for technological advancements to provide deployable solutions for monitoring more wildlife, more accurately, across difficult to monitor spatio-temporal scales (Levin 1992, Estes et al. 2018). Major improvements in sensor miniaturization, imaging resolution, and the development of cheaper unoccupied aircraft systems (UAS or drones) have created new and exciting opportunities for collecting information on the distribution, abundance, and activities of wildlife (Marvin et al. 2016, Hollings et al. 2018). While the scale and data collected using new technologies expands our scientific capabilities, it is not without its own unique challenges. Airborne imaging can collect terabytes of imagery that have historically been manually assessed by researchers to extract information on what species are present in what abundances. While airborne imaging can cover larger expanses more quickly than ground surveys, manual assessment of huge amounts of imagery is still time consuming and error prone (Frederick et al. 2003, Hodgson et al. 2018), creating an obstacle for expanding this approach to wildlife monitoring.

Computer vision algorithms can automate the detection and identification of wildlife in imagery (Weinstein 2018), and are increasingly used to generate counts of individuals and estimates of species abundances for population monitoring (e.g., Groom et al. 2013, Lyons et al. 2019, Yang et al. 2024). Another key aspect of wildlife monitoring – individual-level demographic data such as survival, growth, and reproductive success – is rarely obtained from airborne imaging because these measures typically require repeated observations of an individual over time. Automated approaches to collecting individual-level demographic data from aerial imagery have been explored primarily in plants, which do not move from survey to survey, making automated tracking of individuals across images easier (e.g., Stears et al. 2022, Olsoy et al. 2024). However, when animals become quasi-sedentary for parts of their lives (e.g., nesting), these locations can be used to track unique individuals over time (Sardà-Palomera et al. 2017, Picardi et al. 2020), providing a potential pathway for collecting individual-level demographic data using time series of images.

Nest success - the probability that a nest fledges at least one young - is a critical aspect of avian demography used to monitor conservation and management actions and better understand life history evolution and population dynamics (Frederick and Collopy 1989, Sæther and Bakke 2000, Stephens et al. 2004, Fournier et al. 2021, Lauck et al. 2023). Nest success studies are often limited in the numbers of nests or locations because monitoring nests not only requires frequent - potentially disruptive - visits to track fates of eggs and nestlings (Carney and Sydeman 1999, Champagnon et al. 2019) but finding nests in complex, remote landscapes may be difficult (e.g., Potapov et al. 2013, Junda et al. 2015, McClelland et al. 2016, Afán et al. 2018). Aerial flights have long been used to count nests of waterbirds and colony-nesting birds at large scales (Frederick et al. 1996, Kingsford and Porter 2009), making the use of UAS imagery an obvious next step in monitoring these species. While many studies still count birds and nests by hand (McClelland et al. 2016, Albores-Barajas et al. 2018, Lachman et al. 2020, Dundas et al. 2021, McKellar 2022, Sikora and Marchowski 2023, Francis and Brandis 2024), single-flight approaches combining UAS and computer vision have proven effective for species where landscape complexity is low - allowing every nest to be clearly observed (Afán et al. 2018, Lyons et al. 2019, Yang et al. 2024). Finding active nests to track through time is more challenging for species nesting in complex landscapes or in locations where nesting and roosting birds are commonly intermingled (Sardà-Palomera et al. 2012, Ratcliffe et al. 2015, Francis and Brandis 2024). Most studies using unoccupied aircraft systems (UASs) to assess avian reproductive success use field-identified nests for targeted drone monitoring (Potapov et al. 2013, Weissensteiner et al. 2015, Junda et al. 2015, Gallego and Sarasola 2021, Sikora and Marchowski 2023) or use human photo assessment of broad drone surveys to determine likely nests in images (Sardà-Palomera et al. 2017, Lachman et al. 2020, Valle and Scarton 2022, Francis and Brandis 2024). Locating nests in the field or through human photo assessment in the lab requires substantial time and effort (Descamps et al. 2011, Chabot et al. 2018) that can limit the spatial scales and sample sizes being monitored. Recent studies using time series imagery show that humans can identify nests by finding locations where adult birds are present across multiple flights (Sardà-Palomera et al. 2017). If images can be accurately georeferenced and birds automatically detected, then it should be possible to automate nest detection at landscape scales by identifying locations where birds are repeatedly located over time.

Wading birds in the Everglades are an ideal model system for developing and assessing a time-series approach for detecting nests in large-scale drone imagery. Nesting colonies form every year on tree islands which are monitored aerially using a combination of crewed aircraft and UASs. Birds nest at a variety of positions within the canopy, generating a range of nests from aerially exposed to partially or completely obscured. In addition to airborne monitoring, nests in a subset of colonies are monitored from the ground to collect data on nest success. Here we leverage this combination of an existing airborne monitoring program coupled with on-the-ground monitoring of nests to develop an automated approach for detecting nests and test it against both ground-truthed nest locations and human ability to identify nests from time series imagery.

## Methods

### Study system

The Everglades, located in southern Florida, USA, is a freshwater marsh with higher elevation tree islands embedded in a landscape of deep water sloughs, sawgrass strands, and wetland prairie. Most species of interest are either bright white (White Ibis [*Eudocimus albus*], Great Egrets [*Ardea alba]*, Wood Storks [*Mycteria americana*], Snowy Egrets [*Egretta thula*]), brightly colored (Roseate Spoonbills [*Platalea ajaja*]), or nest conspicuously at the top of the canopy (Great Blue Heron [*Ardea herodias*]). We conducted comparisons of human and algorithm performance using colony-scale weekly imagery from 2020, 2021, and 2022 for 6 colonies (Figure 1). These colonies consisted of several Great Egret dominated colonies (Jerrod, Joule, Vacation, Start Mel), a large mixed species colony (6th Bridge), and a colony with large numbers of Wood Storks (Jetport South). The three years of imagery spanned a range of breeding conditions from relatively poor (2022), to average (2020), to the fourth largest breeding season documented in the study area since 1975 (2021) (Frederick et al. 2021, Ernest and Garner 2022).

**Figure 1.**
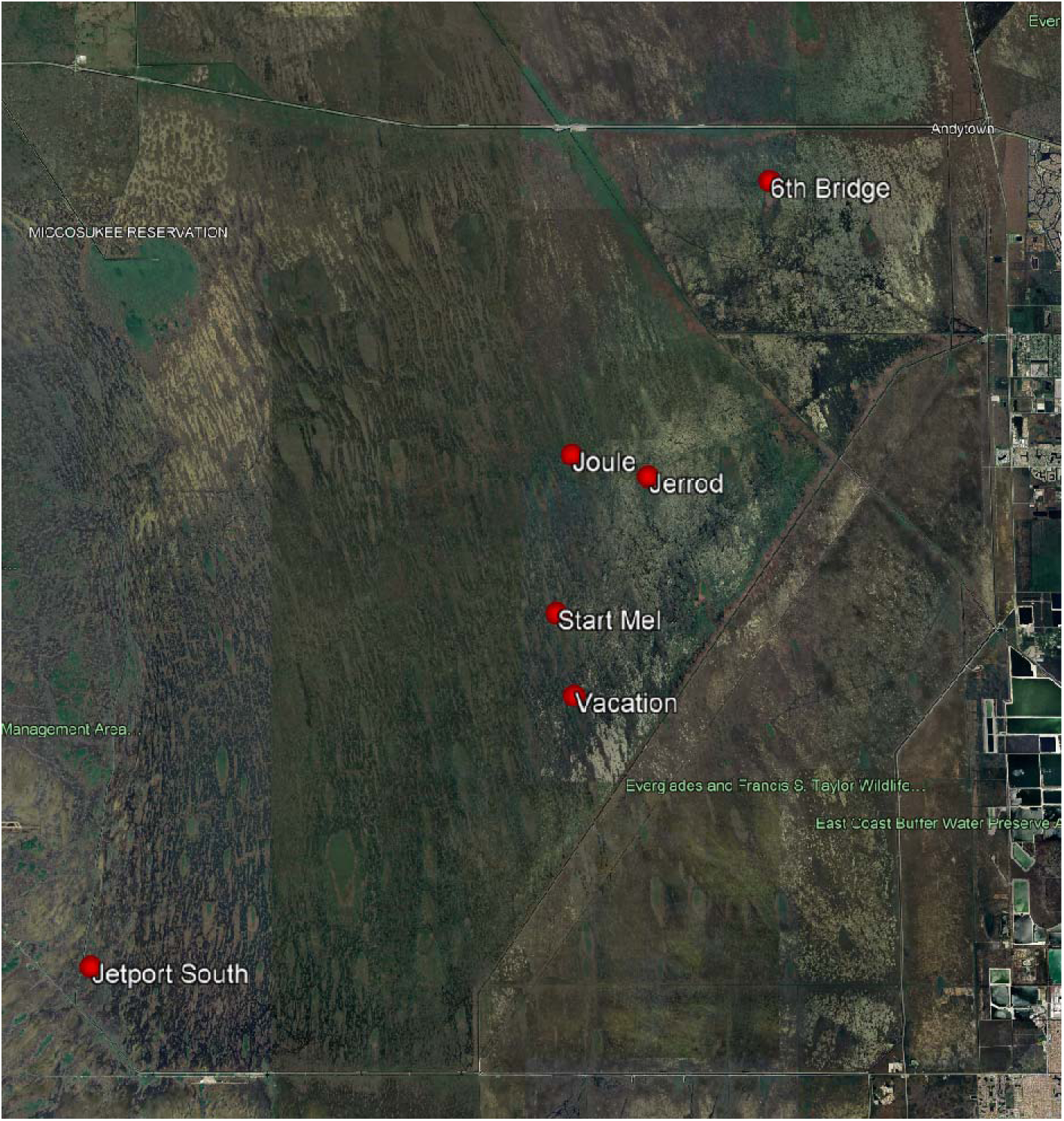
Location of wading bird colonies (red dots) in the Greater Everglades region of southern Florida, USA. Imagery from Google Earth.

### UAS Flights

Weekly flights over colonies roughly coincided with weekly ground nest checks. We used a DJI Inspire II quadcopter equipped with a DJI Zenmuse X7 24 MP camera with a 35 mm equivalent lens and integrated gimbal. Flights were conducted at 250’-300’ AGL which provided a ground sampling distance of ∼1.0 cm. The camera was set at an angle 15 degrees from nadir to provide a slight profile of nesting birds. The DJI Ground Station Pro App mission planner was used to generate repeatable 3D mapping flight plans for each colony that included >75% overlap in all directions to ensure complete colony coverage. To provide accurate positions of birds and nests, each colony had 5-6 ground control points (GCPs) permanently installed around its perimeter.

Each GCP is a 1.5’ x 1.5’ target marked with a checkerboard pattern for easy identification of the exact center point in UAS imagery. The precise location (latitude and longitude) and elevation for each GCP was determined using a Trimble VRS network RTK solution in conjunction with the Florida Permanent Reference Network. Individual images from the UAS were combined for each survey using AgiSoft Metashape software and GCP locations to stitch the overlapping images together into a single georeferenced image of the colony. The GCPs allowed us to generate georeferenced orthomosaics where precise locations of nests could be compared across different flights, which was crucial to our time series approach. Further details of UAS flights and our orthomosaic process can be found in White et al (2024).

### Nest Detection Algorithm

Our nest detection algorithm is based on detecting every bird in each survey using computer vision models and then using instances where a bird is observed repeatedly in the same location across multiple weekly surveys as indicators of nests. To find birds in the orthomosaic image for each survey, we used a RetinaNet-50 object detector that was initially trained on a large compilation of airborne bird remote sensing imagery for general bird detection(Weinstein et al. 2022) and then fine-tuned for the Everglades using tens of thousands of human labeled images from our 2020 and 2021 UAS flights (White et al. 2024). The bird detections from this model are stored as bounding boxes around each bird with a predicted species label. Using these weekly bird locations we then identify birds occurring at the same location through time by sequentially checking each bounding box for intersections with other bounding boxes in the dataset (excluding intersections within the same survey). In cases where a bounding box intersects with multiple bounding boxes in a single survey (due to close proximity between birds) the intersecting bounding box with the greatest overlap (interaction over union; IOU) is automatically chosen as the matching bounding box. Once a bounding box has been assigned to a nest it cannot be assigned to additional nests and is excluded from additional possible matches.

We identify nests based on the number of times a bird occurs at the same location and the continuity of those occurrences. Based on breeding behavior and potential challenges in bird detection due to canopy cover, we designed the ‘bird-bird-bird’ nest identification rule which predicts a nest if overlapping bird detections occurred on three or more different dates at the same location over the breeding season (Figure 2). We chose a minimum of three bird detections because these species continually incubate their eggs for 21-28 days depending on species and young are often confined to the nest for some period thereafter (14-55 days; Coulter et al. 2020, Dumas 2020, Heath et al. 2020, McCrimmon Jr. et al. 2020). We allowed bird detections to be non-consecutive because of detection issues caused by adults flushing from the nest or imperfect visibility (e.g., changes in lighting or wind conditions can obscure birds nesting low in the canopy). An important limitation of the bird-bird-bird approach is that it requires at least three weeks of nest activity. As a result it will miss nests that fail quickly (i.e. are active for less than three weeks), which can be important for determining parameters related to reproductive success (Mayfield 1975). Therefore, we also explored an expanded version of the algorithm that also identified locations where birds were present in two consecutive surveys (i.e. back-to-back weeks) as nests, in addition to the three occurrence bird-bird-bird detections (Figure 2). We refer to this less conservative approach as ‘bird-bird-plus’.

**Figure 2.**
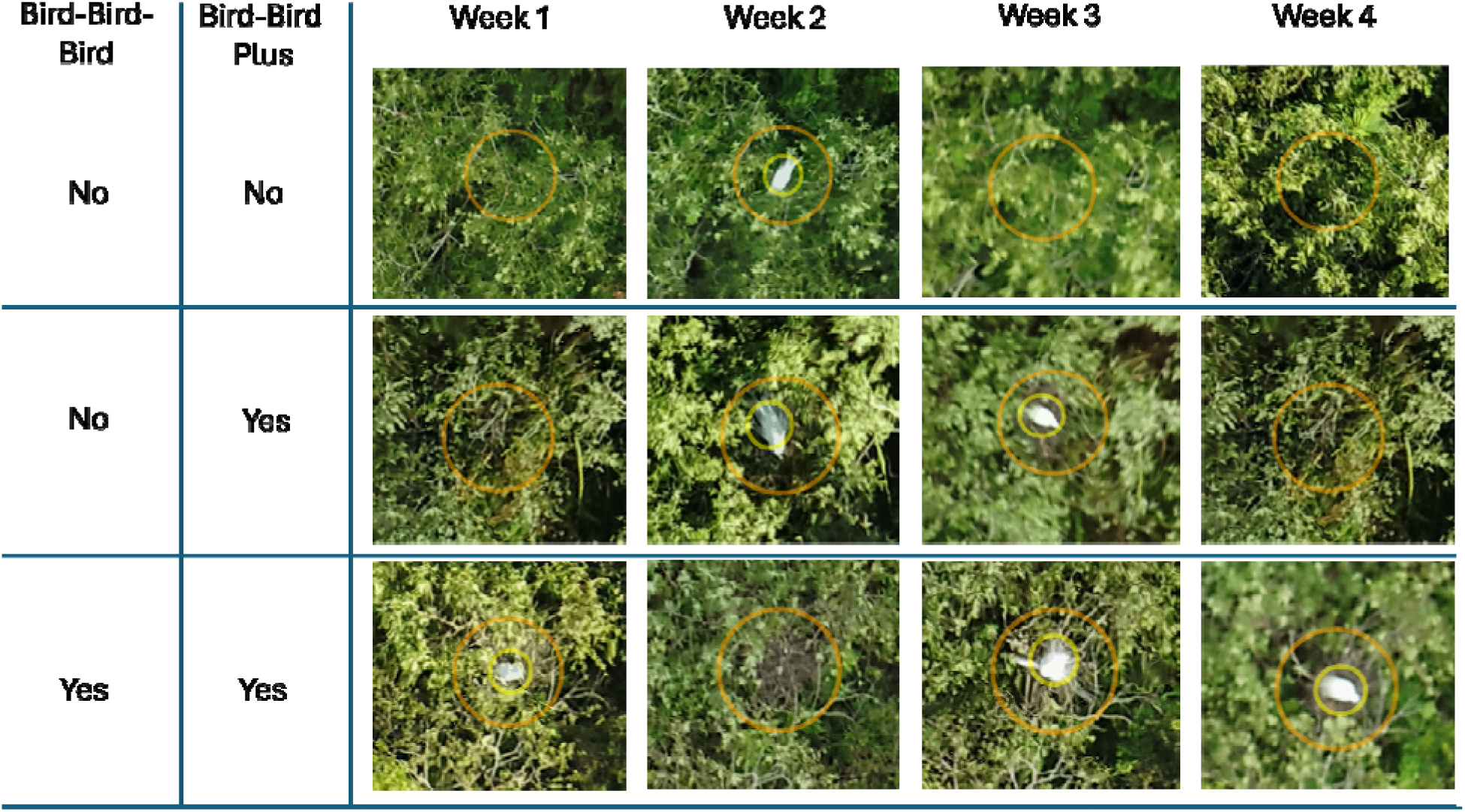
Nest detection examples showing how bird detections are used to predict nest locations using the bird-bird-bird rule (at least three bird detections) and the bird-bird-plus rule (at least three bird detections or two consecutive bird detections). Large orange circle is the target used to assist human assessment of imagery. Small yellow circles indicate automated bird detections which are not seen during human photo assessment.

### Evaluation of Algorithmic Nest Detection

#### Aerial-Only Comparison

The performance of the two algorithms for detecting nest locations was compared to human determination of nest presence using aerial-imagery alone. Algorithmic predictions for nest locations were made at 6 different colonies (Jerrod, Jetport South, 6th Bridge, Vacation, Joule, Start Mel) for the 2020, 2021, and 2022 breeding seasons using bird-bird-bird and bird-bird-plus. Twenty-five algorithmically detected nests were selected for human assessment from each unique year-colony combination. To assess whether nests were missed by the algorithms while avoiding human assessor bias, we provided the human annotator 50 locations to assess: 25 locations where the algorithm had detected a nest and 25 locations where a bird was detected in at least one week, but no nest was algorithmically predicted. This design meant that the human annotator did not know the algorithmic predictions while performing their annotations (Figure 3).

**Figure 3.**
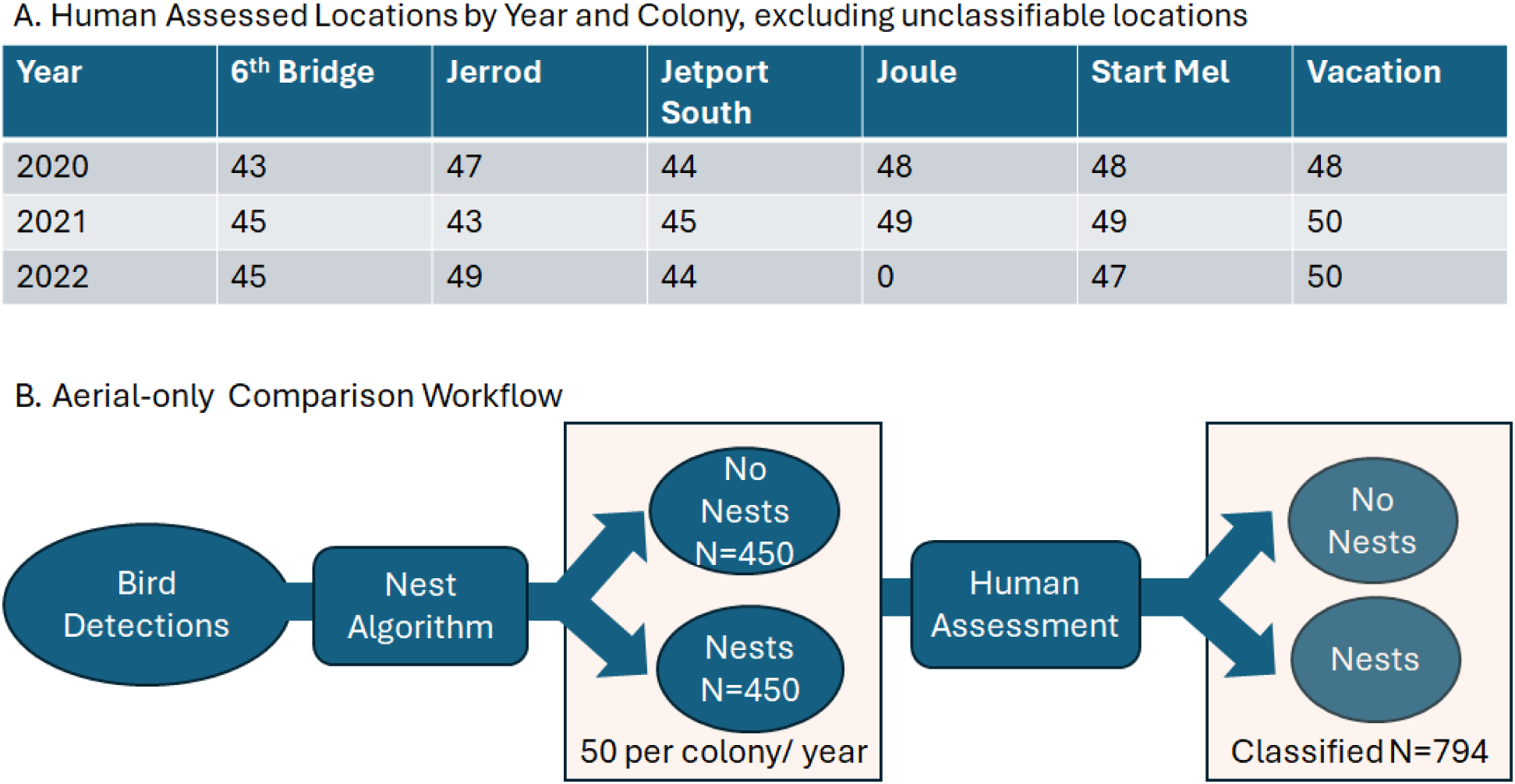
Workflow for airborne image-only comparisons. Bird detections were used by the nest algorithms to predict nest locations. A random subset of locations with and without predicted nests were provided to a human for nest assessment. White boxes indicate data used for performance comparisons.

#### Ground-verified Nest Comparison

We also evaluated algorithmic nest detection and human performance by comparing both to ground-truthed nests based on field surveys. As part of regular long-term monitoring of nesting success, we conducted ground surveys of nests in seven colonies during the breeding season (Jan-June) in 2022. Colonies were chosen for nest surveys based on size, geographic diversity, accessibility by airboat, and species composition to provide a representative sample of nesting success. Nests along established transects were flagged and monitored in weekly visits until the nest either failed or nestlings successfully fledged. For assessment of algorithmic nest detection, a stratified random subset of Great Egret nests in five colonies (Joule, Jerrod, Vacation, Start Mel, and 6th Bridge) were selected to include a variety of heights in the canopy, reflecting the real-world distribution of nests in colonies. When these nests either failed or fledged, GPS coordinates and field notes on nest location were recorded and brightly colored paper was placed in or near the nest (depending upon accessibility) before the next drone flight to help locate the nest in imagery. In an effort to flag as many nests as possible, two Roseate Spoonbill nests were also included in this sample. In addition to flagged Great Egret and Roseate Spoonbill nests, Great Blue Heron nests were also added for assessment based on GPS coordinates and field notes alone because they are easy to locate in drone imagery and pair with specific ground-monitored nests due to their tree-top locations.

Locations of field nests in drone imagery were obtained by a team member involved in the weekly nest checks using QGIS and aided by field notes, handheld GPS waypoints, colored paper placement, and drone imagery collected over the field season. To avoid bias in human performance due to the observer knowing that the images they are assessing represent a field nest, an equal number of random bird locations were also selected for human assessment. Because field monitored nests represent only a small fraction of actual nests in a colony, it is unknown whether randomly selected locations contained nests. Therefore, random locations were only used to reduce human bias and were not used in further analysis (Figure 4). We used the bird-bird-bird algorithm to make predictions for all nest locations in these colonies that could then be compared with the locations of the ground-verified nests to determine if the nest detector located nests at the same locations.

**Figure 4.**
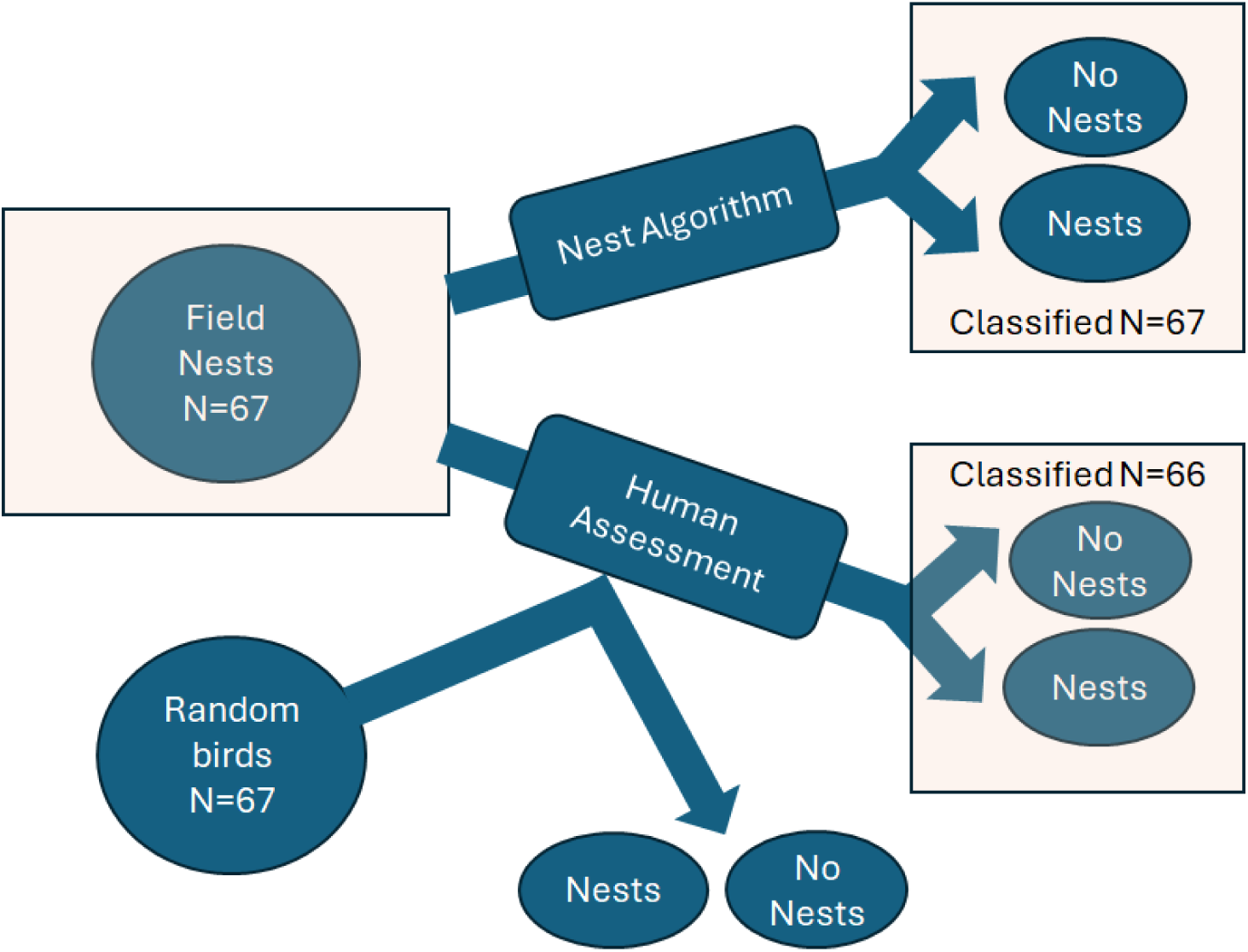
Workflow for Ground-Verified Nest Evaluations. Known nest locations were assessed in the imagery using both the nest detection algorithm and human assessment. Human assessors were also given random bird locations - which might also be unverified nest locations - in order to prevent bias during assessment, but these data were not used in the analysis. Orange boxes indicate data used for performance comparisons.

#### Human Photo Assessment Procedure

To facilitate effective human assessment, we built a Shiny web app to serve the human assessor a time-series of weekly colony-scale orthorectified imagery containing either the 900 pre-selected locations for assessment or the 134 locations that included ground-verified nests. The human observer was trained to identify nests in aerial imagery and had no knowledge of which locations were verified or algorithmically predicted nests. For each assessment location, the app generated a ∼2 m^2^ target circle centered on the sample point (Figure 2) to help the observer focus on the correct location across multiple weekly images. After viewing the time series of images for a location, each assessment location was scored by the annotator as containing a nest, not containing a nest, or undetermined based on: 1) whether a nest was visible; 2) body position; and 3) the frequency and consistency of detections at the location (a designation used primarily for locations partially obscured by canopy cover). Locations were designated as Undetermined if image distortion - a not uncommon issue with georectified images – occurred frequently enough at the location to prevent the human observer from making an assessment. Human photo assessment is not a human-based implementation of the nest detection algorithms and determination of nests by human review can be based on a single image in the time series if that image clearly shows either a nest (with or without a bird) or a bird in a nesting posture.

Post-hoc review of ground-verified nests missed during human review revealed that location uncertainty of nests from field information resulted in a handful of known nests occurring just outside the evaluation target. Therefore, we also report human and algorithmic results using a larger target radius that includes any nests detected touching or immediately adjacent to the ∼2 m^2^ target location. During nest assessment an issue with the georeferencing for the Joule 2022 imagery was also discovered. This issue caused misalignment of locations across weekly images and as a result human and algorithmic assessments were not conducted for this imagery because it was impossible to track a specific location over time. We also discovered bird detection issues with Jetport South in 2022 where tree stumps and branches at the colony edge were detected as birds, probably due to their shape and white coloration in the imagery. While these issues were noted, we continued assessment for this location since issues with algorithmic bird detection – and hence nest detection – are a likely source of error in real-world applications of computational approaches to nest detection. After removing locations with imagery issues (either at the colony or location level) that prevented human assessment, we retained 66 of 67 ground-verified nests and 794 of 900 aerial-only locations.

#### Algorithm Performance

Because true nest locations are not known in the aerial-only comparison, human assessments of the airborne imagery serve as the benchmark for comparison. We calculated two common metrics used in computer vision for assessing model performance: 1) Recall – the fraction of human identified nests that the automated approach also detected; and 2) Precision – the fraction of automated nest detections that were also identified as nests during human photo assessment. Precision and recall were calculated for each year and for each colony. We compared precision and recall values across years and by site, and for the two nest detection algorithms (bird-bird-bird, and bird-bird-plus). For ground-verified nest comparisons we had real ground-truth data, but not all nests in an area were labeled. Therefore, we evaluated performance based on the percent of ground-verified nests detected by the algorithmic approach and by human photo assessment, as well as the percent of human-identified nests that the algorithm also detected.

## Results

### Aerial-only Comparisons

Using human nest assessment as the standard of comparison, we examined colony and year level performance of the bird-bird-bird and bird-bird-plus algorithms for predicting nests in airborne imagery across years and colony types. The bird-bird-bird algorithm showed near human-level performance, finding a majority of the nests identified by the human annotator (recall=0.82) while also generating few false positives (precision = 0.86). Performance varied little across years with both precision and recall values across colonies distributed around the overall mean as well as year-level means closely approaching the overall mean (Figure 5). However, precision was slightly higher than average in 2021 and recall was slightly higher in 2022. There was also a single colony in 2022 with a notably low outlier for precision (Figure 5).

**Figure 5.**
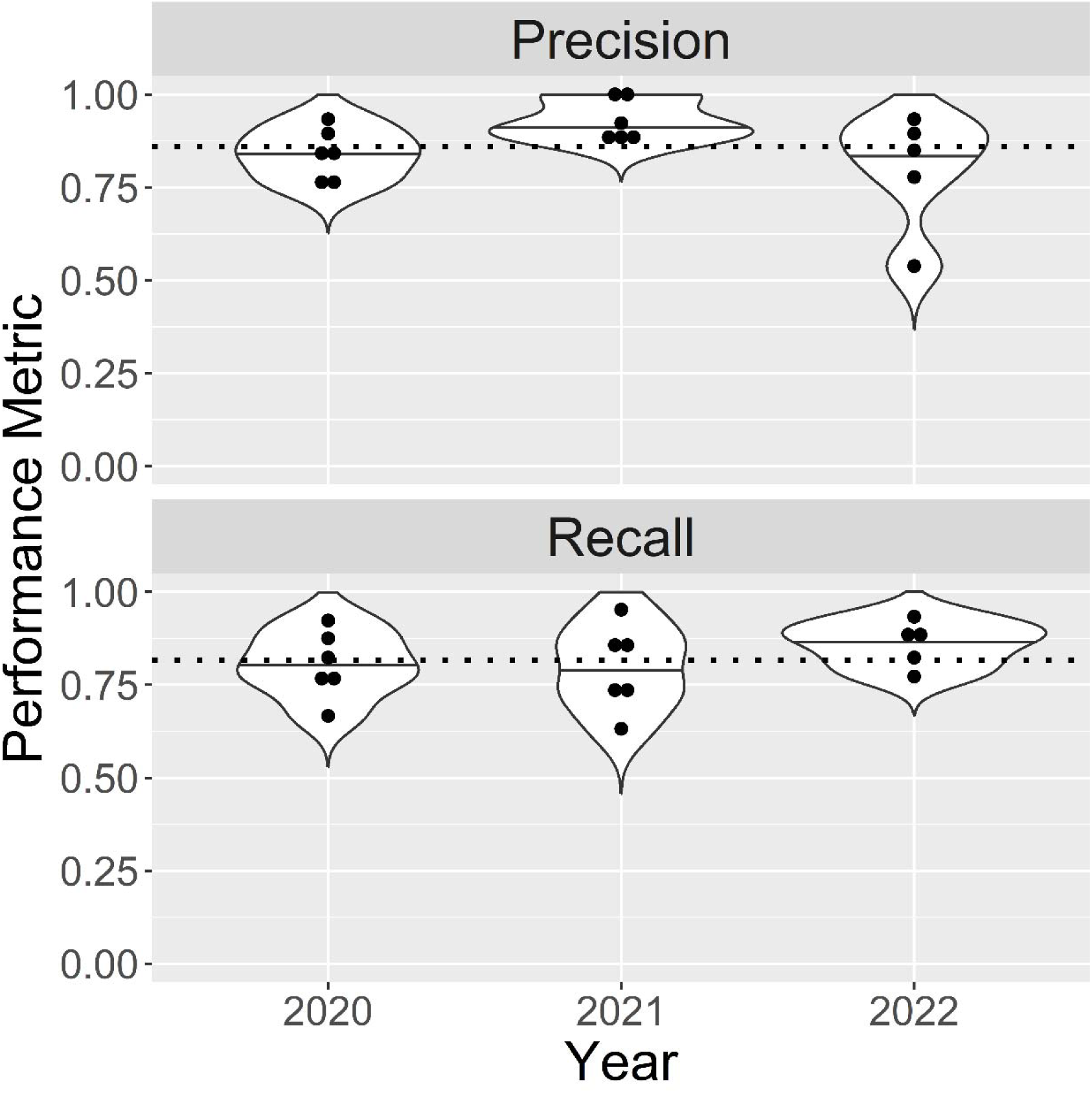
Precision (top panel) and recall (bottom panel) by year for the performance of the bird-bird-bird algorithm compared to human aerial-only labels. Each dot represents the performance measure for an individual colony. The dashed line is the average performance across years and colonies.

There is little evidence from our results for meaningful colony-level differences in precision and recall (Figure 6). The most evident outlier occurs in the precision of the Jetport South colony, but this is driven by a single datapoint, the same datapoint that appeared as an outlier in the 2022 data (Figure 5). This outlier exerts a large influence on the resulting violin plot. The low precision for JetPort South in 2022 was caused by an issue with the bird detection algorithm that erroneously detected silvery gray cypress tree stumps as birds. Because stumps don’t move between images, they were then flagged as potential nests. In general, post-hoc review of differing nest assessments by the algorithm and human annotator indicated a mix of both bird detection issues (including missed bird detections due to obscuring canopy cover) and quick fail nests that did not meet the 3-detection rule.

**Figure 6.**
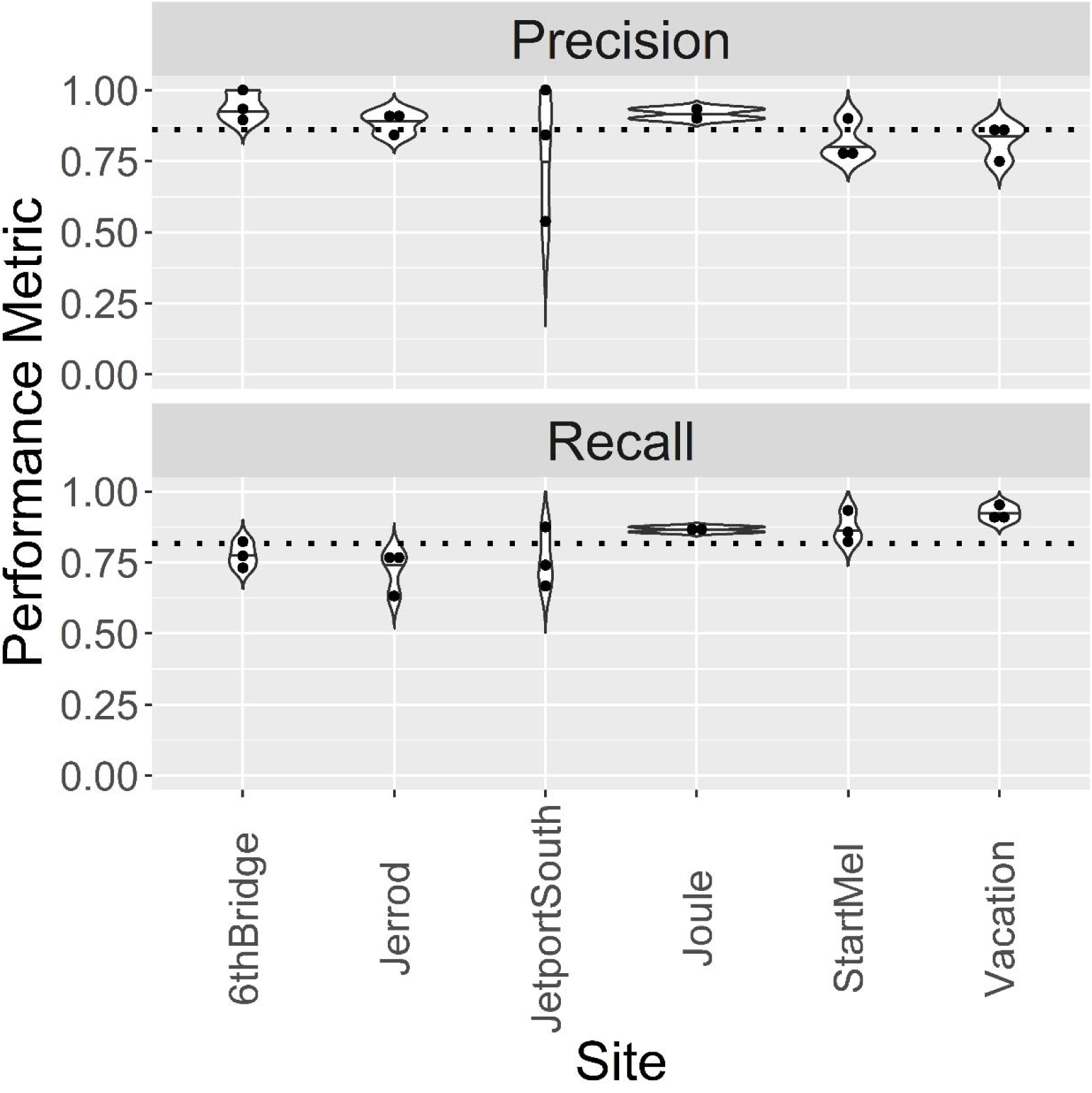
Precision (top panel) and recall (bottom panel) by colony for the performance of the bird-bird-bird algorithm compared to the human aerial-only labels. Each dot represents the performance measure for a single year. The dashed line is the average performance across years and colonies.

### Comparing nest detection rules

We compared precision and recall (for human annotations of imagery) between the bird-bird-bird and bird-bird-plus algorithms to evaluate the potential of bird-bird-plus to locate quick-fail nests. Adding locations with 2-consecutive detections (bird-bird-plus) had little impact on the overall ability to find human-identified nests (recall=0.83 vs. 0.82) but did increase the number of false positives (precision = 0.76 vs. 0.86). Examining results by year and colony (Figure 7) shows these results were generally consistent.

**Figure 7.**
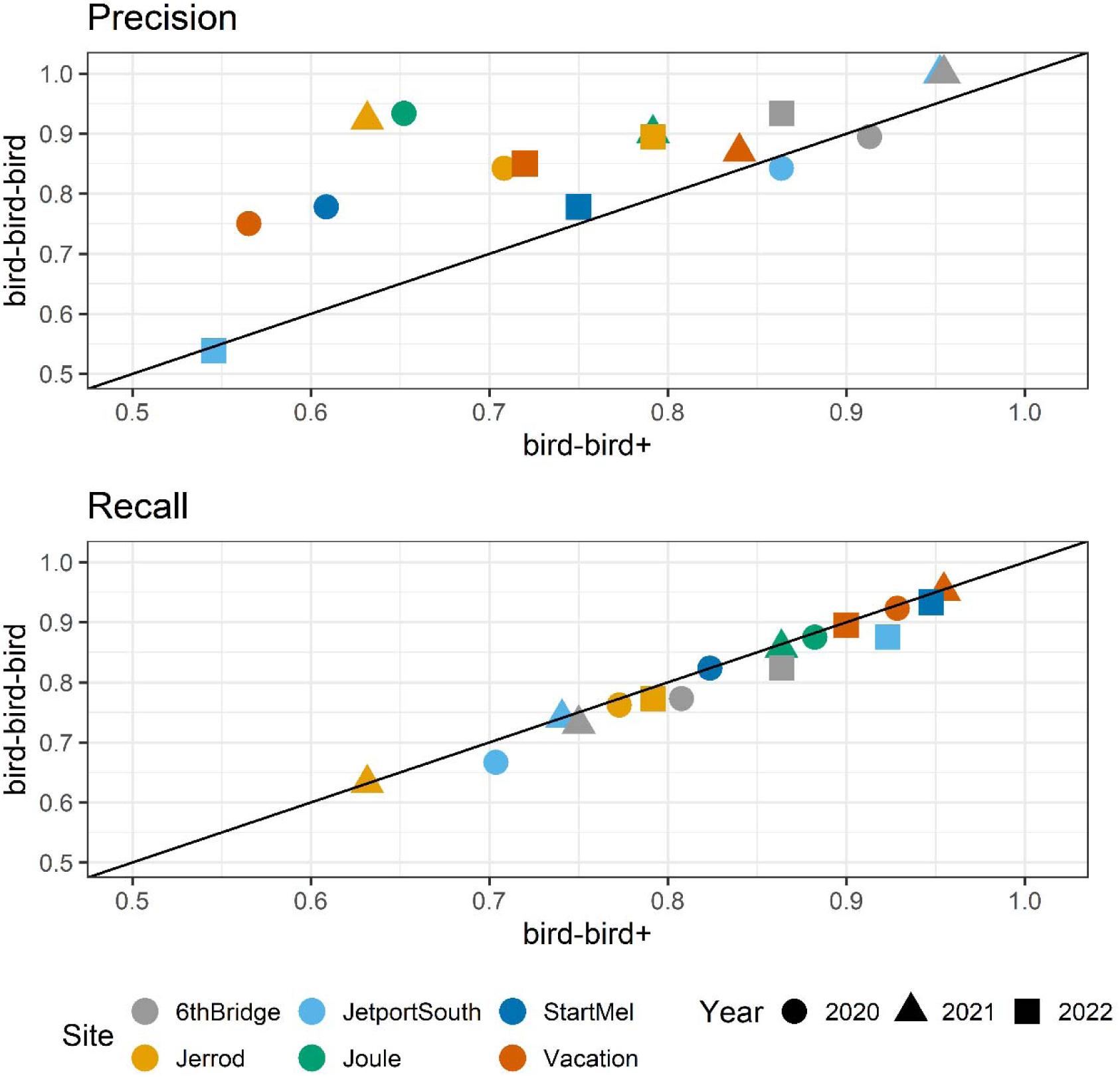
Comparison of bird-bird-bird and bird-bird-plus approaches for precision (top panel) and recall (bottom panel). Each point presents the performance value for a specific colony-year combination. Details in legend.

### Ground-verified nest detection

Automated nest detection using the bird-bird-bird rule found 68% (45 out of 66) of nests identified from ground surveys when only detections strictly within the target area were included. This number increased to 77% (51/66) when detections touching or immediately adjacent to the detection area are also included. In comparison, a human observer using only the UAS imagery to find nests found 74% (49/66) of these nests when only assessing the area strictly within the target area. Including nests detected touching or immediately adjacent to the target area raised this number to 92% (61/66). Given that the inclusion of nests outside the target area also involved post-hoc knowledge which could make it easier to find nests in imagery, these results probably suggest that between ∼74-90% of nests are identifiable by humans from aerial imagery and that the automated approach can find between 83-90% of the nests that humans can locate. This mirrored the results from the aerial-only nest detection where most of the nests identified by human annotation were successfully identified by the algorithm.

## Discussion

Time-series of airborne imagery can be used to quantify important aspects of ecological systems that cannot be assessed from a single flight (e.g., Sardà-Palomera et al. 2017, Olsoy et al. 2024). Using weekly flights throughout a breeding season, we were able to algorithmically identify wading bird nests in imagery with near human-level performance. The bird-bird-bird approach to nest detection (at least three spatially overlapping bird detections across weekly surveys) found most of the nests a human assessor could find in the same imagery (>90%) and performed consistently across years and colony types. Human labelers and algorithmic approaches failed to detect ground-truthed nests at similar rates, with post-hoc assessment suggesting quick fail nests were a frequent cause of missed nests. Canopy cover can also interfere with bird detection - causing birds to be missed by the detector even though portions of them are visible to a human eye. Increased frequency of flights could mitigate these issues by increasing the likelihood of detecting quick fail nests (by ensuring at least three flights before the nest fails) and increasing the chance of catching glimpses of birds under canopy cover. However, increasing the frequency of flights comes with rising labor and travel investments, which need to be weighed relative to other goals such as the desired spatial scale of monitoring. Overall, the performance of our nest detection algorithm suggests automated nest detection can be deployed to locate nests in aerial imagery more quickly and at larger scales than might be feasible with humans alone.

There is no golden rule for automating nest predictions in time series imagery. A previous study using human assessment of nest locations without automation (Sardà-Palomera et al. 2012) used both multiple flights per day (to identify possible nests) and weekly flights (to confirm nest presence). We did not conduct multiple flights per day but instead assessed different rule sets applied to weekly images for making nest predictions. Bird-bird-bird does not require consistent detection at the location, which makes it more flexible and lessens (though does not eliminate) impacts of changes on the ground such as shifts in canopy cover or lighting that alter visibility into the understory through gaps in the canopy. The requirement for three detections also reduces the likelihood of repeat roosting behavior being falsely identified as a nest. However, because we are collecting imagery once/week, it also requires at least 3 weeks of detections. Short duration nests are particularly likely to be missed which can be important for calculating nest survival metrics such as Mayfield nest success (Mayfield 1975). The bird-bird-plus approach was an attempt to detect short-duration nests by also identifying locations with 2-consecutive detections. This approach detected slightly more human-labelled nests (i.e. higher recall) but this marginal improvement came at the cost of a large increase in false positives (i.e., lower precision). False positives can occur when birds roost at the same location across UAS flights – a scenario we often detected late in the season when juveniles had left the nest but had not left the colony. Bird-bird-bird generally outperformed bird-bird-plus for the years we assessed but nesting conditions were generally favorable across the colonies and years used in this study. Under poor nesting conditions, it is possible that bird-bird-plus might outperform bird-bird-bird. Use cases that require detecting as many true nests as possible might still benefit from bird-bird-plus if they have a system for excluding false positives (e.g., using the algorithm to identify possible nest locations for human review). Further assessment under a wider array of nesting conditions is necessary but it seems likely that the best algorithm for detecting nests in time-series imagery will vary depending on the frequency of field observations, environmental conditions, the quality of the breeding conditions, and the use case, thus requiring ongoing human involvement in algorithm selection and assessment.

While a time series approach to nest detection – whether manual or automated – is a promising approach for finding nests across large, complex landscapes (Sardà-Palomera et al. 2017, Lachman et al. 2020), it relies on the ability to detect birds in images and track their locations precisely across multiple images. While conducting this project, we encountered issues with image processing that impacted both bird detection and the consistency of georeferencing of pixels. Georectifying imagery is necessary to precisely compare bird positions across multiple flights but insufficiently accurate georectification prevented us from assessing one colony (Joule) in 2022. While a human may visually track birds across images – though this is difficult without clear landmarks – the current algorithm only knows if a bird was detected at a specific set of coordinates. This is why precisely located ground control points were crucial to this approach, though PPK or RTK enabled drones may also be able to provide similar georeferencing capabilities.

In addition to imperfect georeferencing, locations with complex vertical topography can be distorted during the orthomosaic process. Colonies located on tree islands with a mixed height canopy were blurred by the process, making bird detection more difficult for both automated and human assessment. This issue arose frequently at the Jetport South colony due to the presence of tall cypress trees. Even when orthomosaics are clear of image distortions and consistently georectified, changes in image quality or local conditions can sometimes confuse the bird detector and have downstream impacts on nest detection. Low precision values at Jetport South in 2022 were generated by the bird detector incorrectly identifying small cypress stumps (which appeared white and roundish like a nesting bird) and branches (which would appear white and sinuous like a bird’s neck) as birds. Because plants do not move, they show up repeatedly across flights and become identified as nests, generating false positives. We did not observe this issue with either the 2020 or 2021 imagery for Jetport South, suggesting a subtle shift in 2022 either on the ground or in our image capture. This type of issue can be addressed by improving algorithmic bird detection to avoid false positives (White et al. 2024), but this emphasizes that keeping humans “in the loop” by conducting ongoing quality control assessment is an important component of automated bird and nest detection.

Time-series imagery-based approaches to automate nest detection have a number of benefits for monitoring bird populations. For ecosystems, such as ours, where nest establishment is asynchronous, or where renesting is common, estimating the total number of nests across a season may not be possible from a single airborne survey (Williams et al. 2011). However, using repeated imagery over time, nests can be monitored as they establish and disappear through the course of a breeding season. Because nests can be tracked over time, nest histories can potentially be constructed using the image time series (Sardà-Palomera et al. 2017, Lachman et al. 2020). Automating nest detection will accelerate locating nests across repeat, large scale imagery and can then be used to automatically process this imagery and extract time series images of likely nest locations for human review and assessment. While image-collected nesting data may not be directly equivalent to ground-collected nest data (see Callaghan et al. 2018), it could still be used to provide important information on timing of nest initiation, length of nesting activity at each nest, and other types of reproductive data (e.g., clutch sizes from visible nests) that could then be used to assess and compare populations (Ricklefs and Bloom 1977, Thompson et al. 2001). Further application of airborne time-series to assess nest success (i.e. the probability of a nest fledging at least one chick), especially any automation of that process, will require extensive study, but the potential for obtaining large-scale, large sample size nesting success data across a breeding season has obvious benefits for conservation, management, and development and testing of ecological theory.

Integration of technological advancements into wildlife monitoring is critical for increasing the scale and frequency at which we can assess species and ecosystem dynamics (Estes et al. 2018). Time series approaches in general exhibit great promise for expanding the information collected using technology beyond counts of individuals and species in snapshots of time to more nuanced data on growth, survivorship, and reproduction (Sardà-Palomera et al. 2017, Lachman et al. 2020, Olsoy et al. 2024). Converting the promise of technology into a reality, however, cannot be undertaken by either computational ecologists or wildlife biologists on their own. Our team included computer vision specialists, software engineers, and wildlife ecologists. Combining these very different scientific cultures are necessary to ensure that the data collected is useful, accurate, and can be processed at the scales and frequencies necessary for wildlife monitoring. It will also require financial and institutional support and an increased valuation of the importance of methodological advancements - a perspective that has been, at times, lacking in the question-focused culture of ecology. If these challenges are tackled, it will expand the scale at which individual level demographics can be estimated, significantly improving large scale wildlife monitoring and associated scientific and management activities.

## Acknowledgements

This research was supported by grants from the U.S. Army Corps of Engineers (W912HZ-20-2-0022/3 to S.K.M. Ernest and P. Frederick), the South Florida Water Management District (4500126520 to S.K.M. Ernest and P. Frederick), the Gordon and Betty Moore Foundation’s Data-Driven Discovery Initiative (GBMF4563 to E.P. White), the National Science Foundation (DEB-2326954 to E.P. White and S.K.M. Ernest), and the USDA National Institute of Food and Agriculture via two Hatch Projects (FLA-WEC-005983 to S.K.M. Ernest and FLA-WEC-005944 to E.P. White). We thank Jerry Lorenz for critical logistical assistance that helped maintain our UAS program and Mark Cook for support and feedback on using UAS to monitor wading bird colonies. Finally we thank John Rouse and Cassandra Farley for helping us navigate the complex regulatory landscape surrounding the use of UAS in Florida.

## Author Contributions Statement

*SKME, EPW, LG, BGW, HS, GMY, and PF* conceived of ideas, designed methodology and contributed to interpretation; *LG* collected UAS images and ground data*, EPW, HS,* and *BGW* implemented the bird and nest detection algorithms and Shiny app for image assessment, *SKME* conducted image assessment and analyzed data; *SKME*, *EPW* and *LG* led the writing of the manuscript. All authors contributed critically to the drafts and gave final approval for publication.

## Data Accessibility

All data and code associated with this paper are publicly archived on Zenodo including: 1) the analysis code for generating the figures and statistical results (https://doi.org/10.5281/zenodo.14871558); 2) the code for generating the annotation environments (https://doi.org/10.5281/zenodo.14871820, which is an archive of two specific branches of https://doi.org/10.5281/zenodo.10827317); 3) the code for running the full computer vision workflow including the bird and nest detection algorithms and the resulting data on bird and nest detections (https://zenodo.org/doi/10.5281/zenodo.11127010).

